# The vagus nerve is necessary for the rapid and widespread neuronal activation in the brain following oral administration of psychoactive bacteria

**DOI:** 10.1101/860890

**Authors:** Aadil Bharwani, Christine West, Kevin Champagne-Jorgensen, Karen-Anne McVey Neufeld, Joseph Ruberto, Wolfgang A. Kunze, John Bienenstock, Paul Forsythe

## Abstract

There is accumulating evidence that certain gut microbes modulate brain chemistry and have antidepressant-like behavioural effects. However, it is unclear which brain regions respond to bacteria-derived signals or how signals are transmitted to distinct regions. We investigated the role of the vagus in mediating neuronal activation following oral treatment with *Lactobacillus rhamnosus* (JB-1).

Male Balb/c mice were orally administered a single dose of saline or a live or heat-killed preparation of a physiologically active bacterial strain, *Lactobacillus rhamnosus* (JB-1). 165 minutes later, c-Fos immunoreactivity in the brain was mapped, and mesenteric vagal afferent fibre firing was recorded. Mice also underwent sub-diaphragmatic vagotomy to investigate whether severing the vagus prevented JB-1-induced c-Fos expression. Finally, we examined the ΔFosB response following acute versus chronic bacterial treatment.

While a single exposure to live and heat-killed bacteria altered vagal activity, only live treatment induced rapid neural activation in widespread but distinct brain regions, as assessed by c-Fos expression. Sub-diaphragmatic vagotomy abolished c-Fos immunoreactivity in most, but not all, previously responsive regions. Chronic, but not acute treatment induced a distinct pattern of ΔFosB expression, including in previously unresponsive brain regions.

These data identify that specific brain regions respond rapidly to gut microbes via vagal-dependent and independent pathways, but suggest long-term exposure is required for the chronic brain activity associated with behavioural changes.

## 1. Introduction

The central nervous system (CNS) constantly integrates a rich array of sensory information from peripheral systems. The vast community of intestinal bacteria is one such source of peripheral signals, interacting with the brain through pathways that together comprise the ‘gut-brain axis’ (Collins et al., 2012; Forsythe et al., 2016). Exposure to certain bacterial strains alters anxiety- and depression-like behaviours, along with facets of CNS physiology under normal conditions (Bercik et al., 2011; Bravo et al., 2011; Janik et al., 2016) and in chronic stress (Bharwani et al., 2017). In humans, consumption of a fermented milk product with specific bacterial strains altered the activity and connectivity of brain regions that process affective and sensory stimuli (Tillisch et al., 2013). Similarly, changes in brain activity have been measured by functional magnetic resonance after four weeks of probiotic treatment (Bagga et al., 2018). While such observations have added to a growing literature (Forsythe et al., 2016; Fülling et al., 2019), there remains much work to be done to understand the mechanisms underlying the detection, transmission, and processing of bacteria-derived signals in the brain. Although it is evident that the gut microbiota and certain exogenous strains influence behaviour and neural function, the precise regions in the brain that are recruited in response to bacterial signals, as well as the changes in the response pattern following acute versus chronic exposure to such signals, remain unknown. Additionally, while several pathways have been proposed to mediate such interactions, including neural, immune, and humoural signals, it is unclear whether bacteria recruit multiple pathways, and whether these transmit information to distinct regions of the brain.

c-Fos and ΔFosB are part of the Fos-Jun family of transcription factors that are expressed in response to various stimuli and bind to sites on gene promoters (Nestler et al., 2001a; Mcclung et al., 2005; Nestler, 2015). *c-Fos* is an immediate early gene marker of neuronal activity (Morgan and Curran, 1991). c-Fos protein expression is closely linked temporally with *c-Fos* transcription (Sharp et al., 1991), is maximal at 1-2 hours following a stimulus, and degrades within 6 hours (Sharp et al., 1991; Jung et al., 2014), thus providing a useful indicator of an acute neuronal response. ΔFosB is a truncated splice variant of FosB, a group of proteins that peak at 6 hours following a stimulus (Hope et al., 1994; Chen et al., 1997). However, unlike the full-length protein and other splice variants, the 35-37kD ΔFosB isoform possesses unique properties that render it stable and allow it to accumulate at high levels following repeated exposure to the stimulus. ΔFosB has been studied in several brain regions for its role in mediating behavioural changes through persistent neural adaptations (Nestler, 2015).

In the present study, we examined for evidence of neuronal activity in the brain shortly after oral administration of a specific bacterial strain, *Lactobacillus rhamnosus* (JB-1), previously demonstrated to attenuate anxiety and depression-like behaviours in mice (Bravo et al., 2011; Bharwani et al., 2017). We observed increased c-Fos expression in distributed brain regions within 165 minutes. While both the live and heat-killed bacteria increased firing of vagal fibres, the effects on c-Fos expression in the CNS were mostly limited to the live strain. Sub-diaphragmatic vagotomy abolished c-Fos expression in most but not all regions, further underscoring the role of the vagus and indicating the recruitment of additional, vagal-independent pathways. Finally, only chronic bacterial treatment induced ΔFosB expression in distinct brain regions and changes in behaviour.

## 2. Methods and Materials

### 2.1. Animals

Adult male BALB/c (BALB/cAnNCrl) mice aged 6-8 weeks were obtained from Charles River (Montreal, QC, Canada) and allowed to habituate to the animal facility for at least 1 week. Mice were maintained on a 12 h light/dark cycle (lights on at 5am) with *ad libitum* access to food and water. Prior to experiments, mice were habituated and handled by the researcher. All experiments were conducted in accordance with the guidelines of the Canadian Council on Animal Care and were approved by McMaster University’s Animal Research Ethics Board.

### 2.2. Preparation and treatment with bacteria

*Lactobacillus rhamnosus* (JB-1) bacteria were prepared as described previously (Bravo et al., 2011). Heat-killed bacteria were prepared as previously described (Kamiya et al., 2006; Mao et al., 2013), by heating 10^10^ CFU aliquots of viable bacteria for 20 minutes at 80°C. No bacterial growth was detected after 72 hours under anaerobic conditions at 37°C.

Animals were orally gavaged with 200μl of either phosphate buffered saline (PBS) alone or 2×10^9^ colony forming units (CFU) of live or heat-killed *Lactobacillus rhamnosus* (JB-1) washed and re-suspended in PBS. For experiments involving acute treatment, animals were gavaged with a single dose ∼165 minutes prior to the administration of anesthesia for transcardial perfusions. For experiments involving chronic treatment, animals were gavaged daily over a period of 14 days, with the final treatment occurring 20-24 hours prior to perfusions. This time-point was chosen in an effort to ensure complete degradation of stimulus-induced FosB, which occurs within 18 hours, and ensure labeling of only the stable ΔFosB isoform (Perrotti et al., 2004; McHenry et al., 2016). Control groups in this experiment were treated with oral PBS for two weeks or oral PBS for 13 days followed by a single dose of the bacteria on the final day (acute JB-1).

### 2.3. Vagotomy

Animals underwent surgery for sub-diaphragmatic vagotomy as previously described (van der Kleij et al., 2008; Bravo et al., 2011). Briefly, mice were anesthetized using isoflurane. An upper midline laparotomy was used to visualize the stomach and lower esophagus, and the intestine was retracted after incising the abdominal wall along the ventral midline. The left lateral lobe of the liver was retracted and a ligature was placed at the gastroesophageal junction to enable retraction and expose both vagal trunks, which were dissected. All neural and connective tissue surrounding the esophagus below the diaphragm were removed to transect all small vagal branches. Animals were allowed to recover for 14 days prior to treatment and perfusion. Sham vagotomy was also performed on surgical control animals.

### 2.4. Mesenteric nerve recording

Tissue was prepared from mice as previously described (McVey Neufeld et al., 2013). Briefly, 2 cm fresh segments of jejunum were placed in a 2 mL recording dish lined with Sylgard and filled with Krebs. The oral and anal ends were cannulated and flushed with plastic tubing, and the mesentery was pinned to isolate nerve bundles by microdissection. The serosal compartment was separately perfused with prewarmed Krebs/nicardipine (3 *µ*M). The nerve bundle was gently sucked into a glass pipette with an attached electrode and extracellular multiunit nerve recordings were made using a Multi-Clamp 700B amplifier and Digidata 1440A signal converter (Molecular Devices). For intraluminal administration experiments, baseline recordings were collected for 20 minutes with luminal perfusion of the Krebs prior to adding treatments at equivalent doses to the respective oral treatments identified in methods. For *in vivo* administration experiments, recordings were made from mice that were administered a single dose of saline, live, or heat-killed JB-1, as described above. Electrical signals were bandpass-filtered at 2 kHz and sampled at 20 kHz. Single units representing discharge from individual single vagal fibres were discriminated and identified by their unique spike waveform shape and amplitude (Rong et al., 2004; Perez-Burgos et al., 2013) using Dataview computer software (Heitler, 2007).

### 2.5. Tail Suspension Test

165 minutes following administration of a single dose of treatment or 1 day following a 14-day course of oral treatments, animals were tested for depression-like behaviour with the tail suspension test (TST). Mice were transferred from the housing room to the behavioural testing room and allowed to habituate for 30 minutes. Following habituation, mice were suspended by the tail using 17cm laboratory tape (Can et al., 2012) from a suspension bar. 2cm of the tape was affixed to the mouse tail and the remainder of the tape used for suspension. Animals were suspended for a total of 6 minutes. Behaviour was video recorded and scored by a blinded observer. Freezing behaviour was measured and calculated as a percentage of the total time suspended.

### 2.6. Immunohistochemistry

Mice were anaesthetized with a mixture of ketamine/xylazine solution and perfused with ice-cold heparinized PBS, followed by 4% paraformaldehyde (PFA). Brains were extracted and placed overnight in 4% PFA at 4°C before being rinsed with PBS and stored in solutions of 10% then 30% sucrose in PBS until the brains sank. Fixed brains were snap frozen with isopentane and stored at - 80°C until sectioning. We obtained 50 *µ*m coronal at −20°C and stored the slices in PBS at 4°C. Floating sections were washed 3x in PBS and blocked for 2h in 0.2% Triton-X 100/0.01 M PBS (PBS-T) containing 5% (v/v) normal goat serum. We incubated sections overnight at 4°C with a primary c-Fos antibody in blocking buffer (1:5000, rabbit anti-c-Fos, Synaptic Systems, 226 003), a primary FosB/ΔFosB antibody (1:2000, rabbit anti-FosB, Abcam, ab184938), or a primary TPH2 antibody (1:500, rabbit anti-TPH2, Abcam, ab184505). Unlike the stable ΔFosB isoform, stimulus-induced FosB is degraded by 18 hr. Thus, all cells labeled with the pan-FosB antibody were considered to reflect ΔFosB (Perrotti et al., 2004; McHenry et al., 2016). Sections were washed 3x in PBS-T, then incubated with AlexaFluor 488-conjugated goat anti-rabbit secondary antibody (1:300, Thermo Fisher Scientific, A-11034) for 1h at room temperature. Sections were washed 3x in PBS-T and mounted using Vectashield with DAPI (Vector Labs).

### 2.7 Fos analysis

Sections containing the regions of interest (ROI) were imaged on an Olympus VS120 Virtual Slide microscope or a Zeiss AxioImager Z1 microscope (AxioCam MRm3) at the Research Institute of St. Joe’s, Hamilton. All groups were imaged using identical scanning parameters (gain, exposure time) and laser power. Using ImageJ Fiji (Schindelin et al., 2012; Rueden et al., 2017), images were converted to 16-bit, background was subtracted, and ROIs were traced using recognizable landmarks from The Mouse Brain Atlas, fourth edition (Paxinos and Franklin, 2004): nucleus accumbens (NAc; approximate bregma AP +0.97 mm), bed nucleus of the stria terminalis (BNST; approximate bregma AP +0.01 mm), paraventricular nucleus of the hypothalamus (PVH; approximate bregma AP −0.82 mm), paraventricular nucleus of the thalamus (PVT; approximate bregma AP −1.43 mm), dorsal hippocampus (dHpc; approximate bregma AP −1.67 mm), basolateral and central amygdala (BLA, CeA; approximate bregma AP −1.67 mm), ventral hippocampus (vHpc; approximate bregma AP −3.15 mm), periaqueductal grey (PAG; approximate bregma AP −4.23 mm), rostral dorsal raphe nucleus (DRN; approximate bregma AP −4.23 mm), locus coeruleus (LC; approximate bregma AP −5.41 mm), nucleus tractus solitarius (NTS; approximate bregma AP −6.59 mm). Fos immunoreactivity was quantified using automated thresholding and particle analysis using consistent parameters. Fos^+^ cell nuclei density was calculated by the number of Fos^+^ cells divided by the ROI area (cells/mm^2^).

### 2.8 Statistical analysis

Sample sizes were initially determined based on prior work employing similar methodology; no formal techniques were used to predetermine sample sizes. Certain brain areas could not be quantified for specific subjects due to damage incurred during extraction and processing (please consult figure legend for the *n* of each analysed region). In all experiments, animals were randomly assigned to treatment and surgery groups. Data were analyzed in GraphPad Prism 6 using a two-tailed student’s *t*-test (or a Mann-Whitney U test when assumptions of a normal distribution were violated), and univariate ANOVA (or Kruskal-Wallis test when assumptions for equality of group variances were violated) followed by Tukey’s or Dunn’s multiple comparisons test when significant main effects were observed. Data are shown as mean ± standard error except where otherwise indicated. Statistical significance was set at *p* < 0.05.

## 3. Results

### 3.1. A single orally administered dose of live bacteria induces c-Fos immunoreactivity throughout the brain

Given multiple converging lines of evidence demonstrating the influence of gut-brain signalling (Collins et al., 2012; Cryan and Dinan, 2012; Forsythe et al., 2016), we interrogated which brain regions were recruited by acute oral exposure to *Lactobacillus rhamnosus* (JB-1)—a strain previously demonstrated to be physiologically active (Bravo et al., 2011; Perez-Burgos et al., 2013, 2014; Bharwani et al., 2017). Seven of 16 regions exhibited robust c-Fos expression only in response to the live but not heat-killed bacteria or saline within 165 minutes following oral administration. This included the basolateral and central amygdala (Fig. 1A: BLA, *F*_(2,18)_= 6.616, *p*= 0.007, Tukey post-hoc live vs heat-killed, *p* < 0.05; Fig.1B: CeA, *F*_(2,18)_= 16.5, *p*< 0.0001, Tukey post-hoc live vs heat-killed, *p* < 0.001). In the ventral hippocampus, the vCA1 showed significantly elevated c-Fos expression in response to the live bacteria only (Fig. 1C: *F*_(2,18)_= 5.769, *p*= 0.0116, Tukey post-hoc, *p* < 0.05). In the vCA3, relative to saline treatment, there was a significant increase in c-Fos^+^ nuclei density in mice that were administered the live bacteria (Fig. 1D: *F*_(2,18)_= 4.756, *p*= 0.022, Tukey post-hoc, *p* < 0.05). Examination of the periaqueductal grey and dorsal raphe nucleus revealed a significant effect of treatment (Fig. 1E: PAG, *F*_(2,18)_= 12.33, *p*= 0.0004; Fig. 1F-H: DRN, *F*_(2,18)_= 9.602, *p*= 0.0015). Both regions showed elevated c-Fos expression in response to administration of the live bacteria only. Similarly, there was a main effect of treatment in the locus coeruleus (Fig. 1I: LC, *F*_(2,18)_= 9.542, *p*= 0.0015), with mice that were administered the live bacteria showing significantly greater c-Fos expression. Only the paraventricular nucleus of the thalamus exhibited a significant c-Fos response to administration of live and heat-killed JB-1 (Fig. 1J: PVT, *F*_(2,18)_= 5.164, *p*= 0.0169; Tukey post-hoc, live and heat-killed JB-1 vs saline, *p*< 0.05).

**Figure 1.**
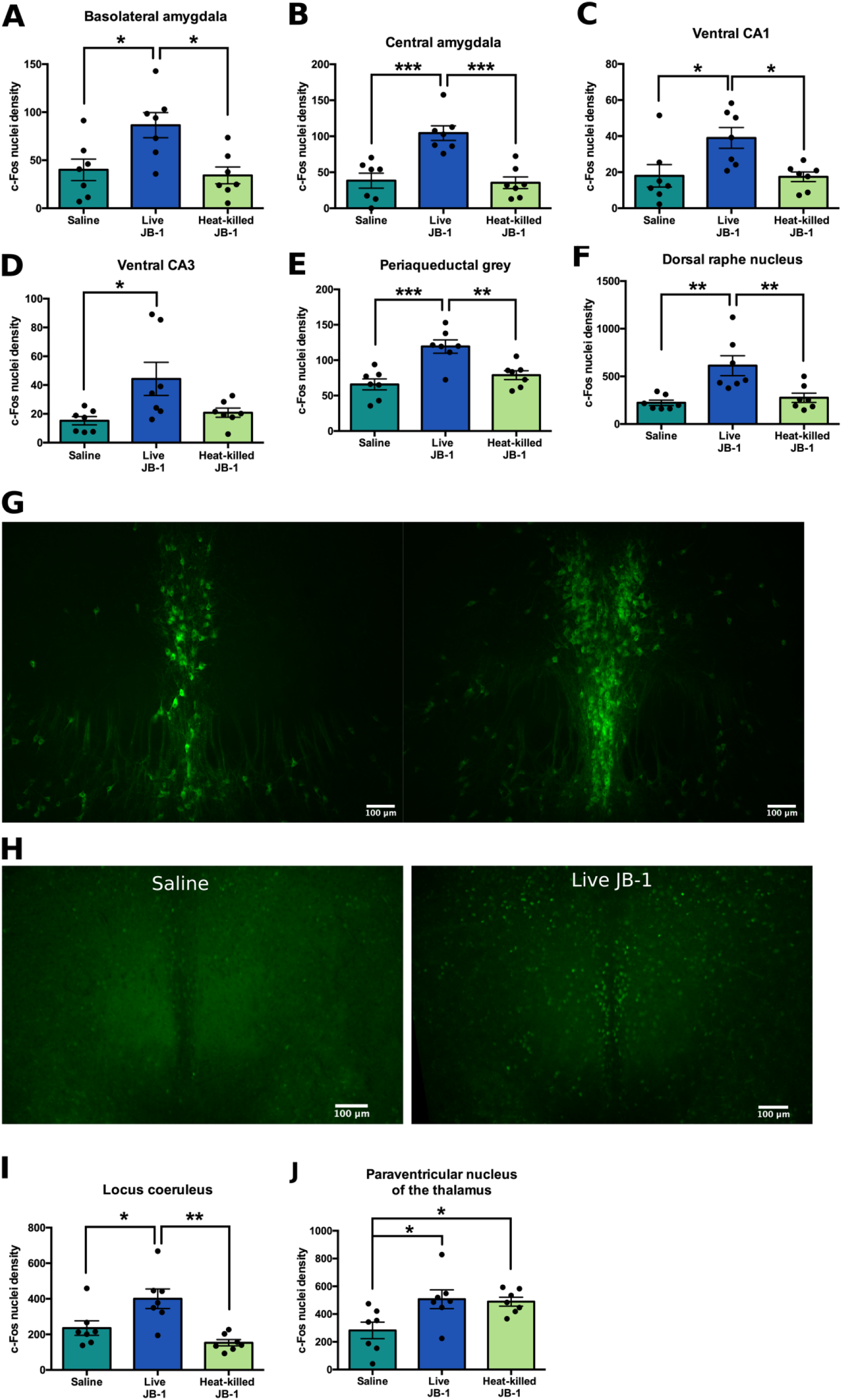
A single oral dose of live bacteria causes an increase in c-Fos expression in specific regions. **(A-F, I-J)** c-Fos^+^ cell density (cells/mm^2^), quantified 165 minutes following administration of saline, live JB-1, or heat-killed JB-1 (n=7 for all groups). **(G)** Images of tryptophan hydroxylase 2 staining in the DRN at locations representative of c-Fos quantification. **(H)** Representative images of c-Fos staining in the DRN * p<0.05, ** p<0.01, ***p<0.001.

In contrast, eight of the 16 regions exhibited no significant c-Fos response to either the live or heat-killed bacteria at 165 minutes following treatment. This included the nucleus accumbens (Fig. 2A: NAc, *H*_(2,18)_= 4.275, *p*= 0.1183); bed nucleus of the stria terminalis (Fig. 2B: BNST, *F*_(2,18)_= 1.832, *p*= 0.1888); paraventricular nucleus of the hypothalamus (Fig. 2C: PVH, *F*_(2,18)_= 0.0830, *p*= 0.9207); dorsal CA1, CA3, and dentate gyrus (Fig. 2D: dCA1, no c-Fos detected; Fig. 2E: dCA3, *F*_(2,18)_= 0.0021, *p*= 0.9979; Fig. 2F: dDG, *F*_(2,18)_= 1.073, *p*= 0.3627); ventral dentate gyrus (Fig. 2G: vDG, *F*_(2,18)_= 0.8120, *p*= 0.4596); and nucleus tractus solitarius (Fig. 2H: NTS, *F*_(2,17)_= 1.405, *p*= 0.2724). Taken together, these data demonstrate that oral administration of a specific bacterial strain is sufficient to activate multiple brain regions within 165 minutes, and indicate the specific nature of this signalling to the live bacteria only.

**Figure 2.**
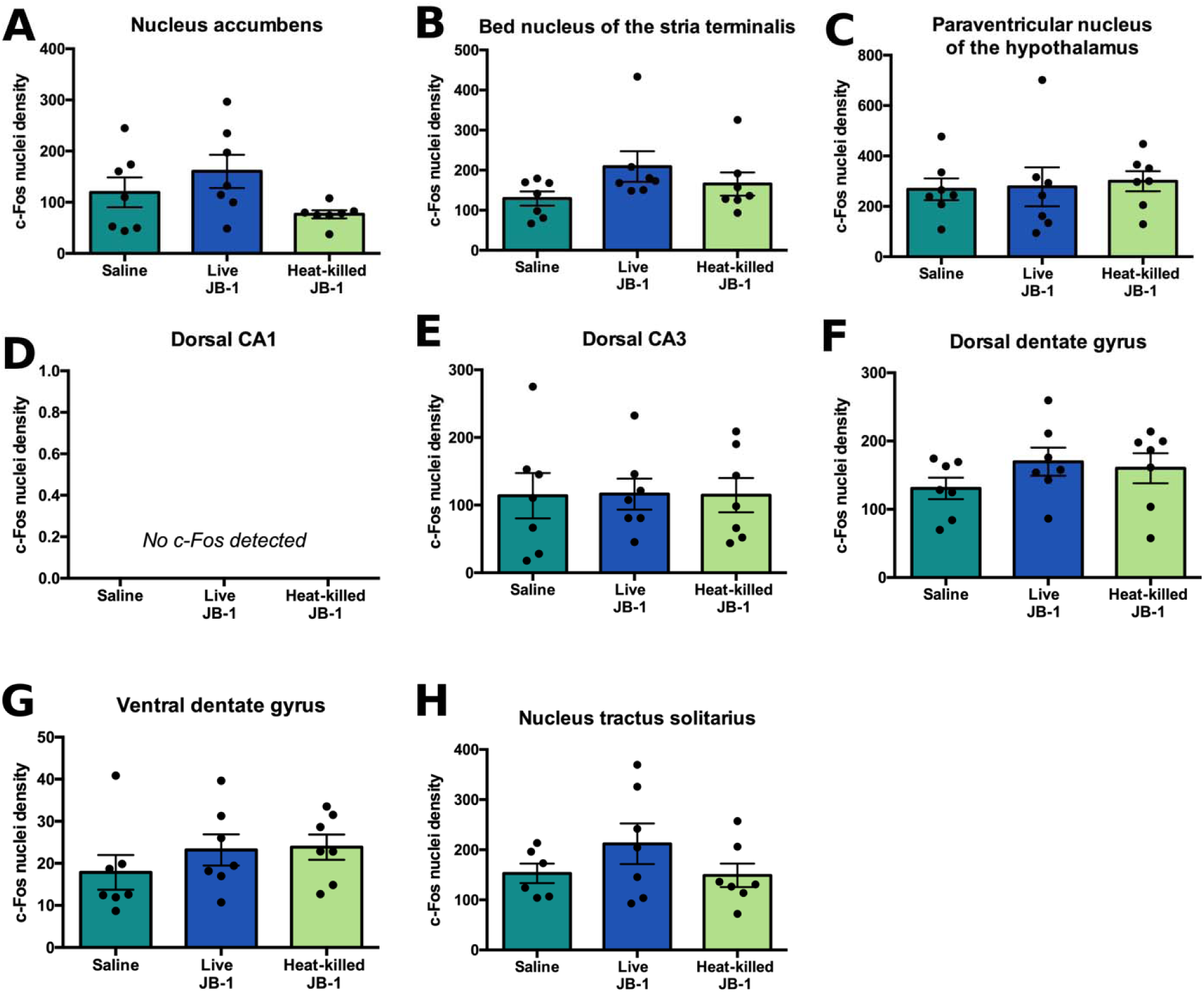
A single oral dose of live bacteria does not affect c-Fos expression in certain regions. **(A-H)** c-Fos^+^ cell density (cells/mm^2^), quantified 165 minutes following administration of saline, live JB-1, or heat-killed JB-1. There were no significant differences in c-Fos levels between treatment groups. n=7 for all groups, with the exception of the NTS of the saline control group (n=6) **(H)**, due to damage incurred during dissection and tissue processing.

### 3.2. Live and heat-killed bacteria facilitate firing of vagal afferent fibres

We have previously demonstrated that live JB-1 increases the firing of vagal afferents via a nicotinic synapse (Perez-Burgos et al., 2013, 2014), and that an intact vagus nerve is necessary for the effects of chronic JB-1 treatment (Bravo et al., 2011). Given that heat-killed JB-1 increased neuronal activity in the PVT (Fig. 1G), we wondered whether the heat-killed bacteria also induced a vagal response. We recorded afferent vagal fibre activity from the mesenteric nerve bundle after adding either the live or heat-killed bacteria directly into the gut at equivalent doses to the oral treatment. Exposure to either treatment significantly decreased the interval between spikes, indicating increased vagal firing (Fig. 3A: live JB-1, t_(31)_=4.606, *p*< 0.0001; Fig. 3B: heat-killed JB-1, t_(29)_= 4.173, *p*= 0.0002). Comparison of treatments showed no significant difference in the change in interspike interval (Fig. 3C; *U*= 420, *p*= 0.6671); however, there was a significant difference in the variance (*F*_(28,30)_= 4.507, *p*= 0.0001). Next, we measured vagal activity 165 minutes after mice were orally administered treatments. Only vagal fibres in the live bacterial treatment group exhibited lower interspike interval 165 minutes following treatment (Fig. 3D; *F*_(2,86)_= 6.113, *p*= 0.0033; Tukey’s multiple comparison test, saline versus live JB-1 groups, *p*< 0.01). These data suggest that while both live and heat-killed JB-1 facilitate vagal firing, there may be differences in the pattern and responsivity of individual vagal fibres, which may account for differences in the c-Fos response.

**Figure 3.**
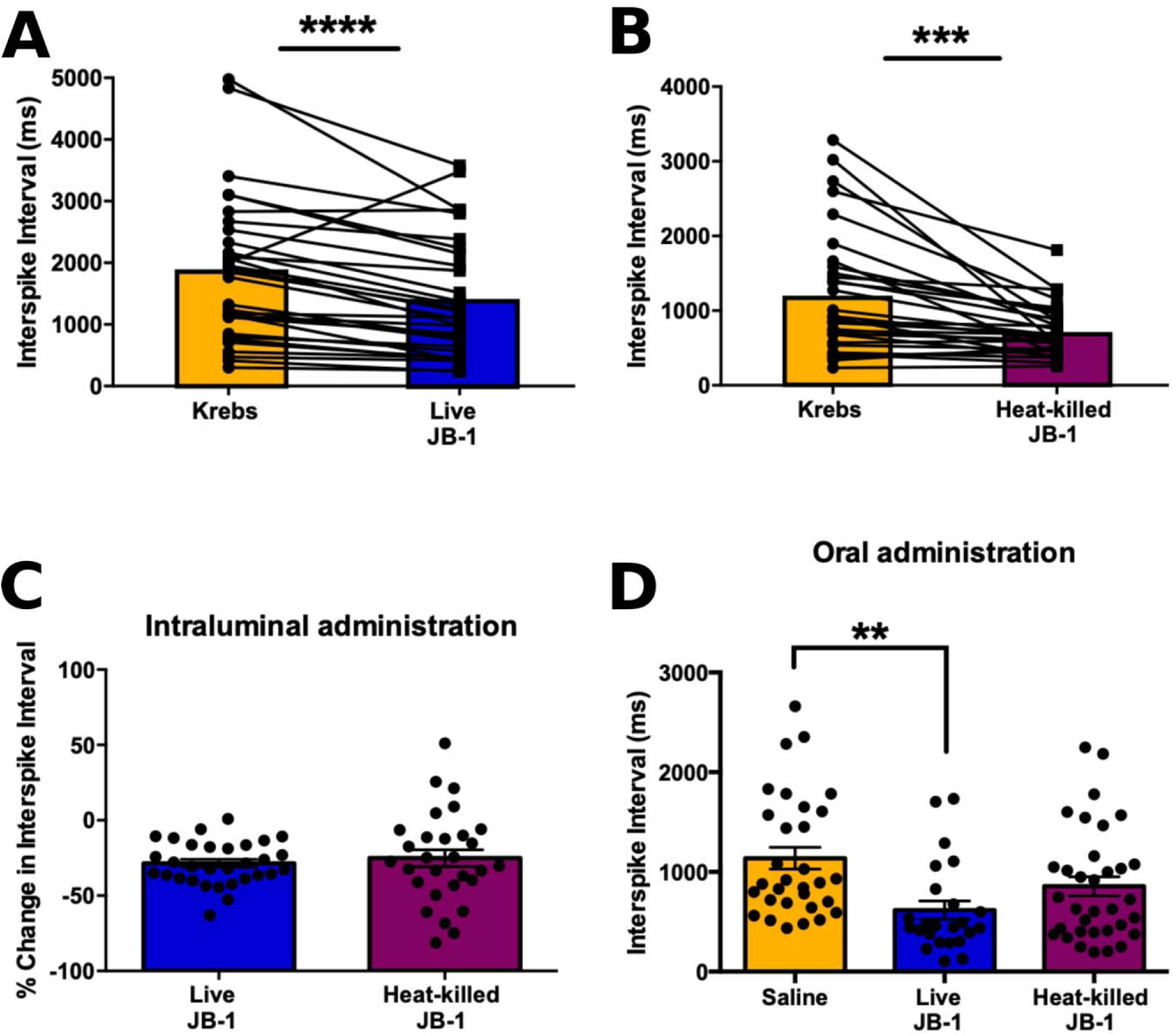
Effect of bacteria exposure on mesenteric vagal afferent fibre activity. **(A & B)** Change in time interval between vagal spike firing following direct, acute exposure of the jejunal lumen to **(A)** the live (n=5 animals, 32 fibres) or **(B)** heat-killed bacteria (n=5 animals, 30 fibers) at equivalent dose to the oral treatment. **(C)** Percentage change in time interval between vagal spike firing in jejunal segments directly exposed to live versus heat-killed bacteria treatment. **(D)** Change in time interval between vagal spike firing in jejunal segments, measured 165 minutes following oral administration of saline (n=5 animals, 31 fibers), live (n=5 animals, 24 fibers), or heat-killed bacteria (n= 5 animals, 34 fibers). * p<0.05, ** p<0.01, ***p<0.001.

### 3.3. Sub-diaphragmatic vagotomy prevents c-Fos expression in certain regions

While we have previously shown that the vagus nerve is critical for the neural and behavioural effects of JB-1 (Bravo et al., 2011), several pathways have been proposed to mediate gut-brain signalling (Collins et al., 2012) and it is unclear whether any of these are recruited by JB-1 in addition to the vagus. Thus, we examined whether severing the vagus abolished JB-1-induced c-Fos immunoreactivity in the aforementioned regions (Fig. 1). Mice that underwent a sub-diaphragmatic vagotomy (Vx) or sham surgery were orally administered the live bacteria or saline to measure c-Fos expression in the brain after 165 minutes. Vx prevented the effects of JB-1 on c-Fos expression in the BLA (Fig. 4A; *F*_(2,18)_= 5.785, *p*= 0.0115; Tukey’s post-hoc, JB-1 in sham versus Vx groups, *p*< 0.05) and the CeA (Fig. 4B; *F*_(2,18)_= 9.643, *p*= 0.0014; Tukey’s post-hoc, JB-1 in sham versus Vx groups, *p*< 0.05). In the PVT, Vx animals treated with the live bacteria failed to show an increase in c-Fos (Fig. 4C; *F*_(2,18)_= 8.062, *p*= 0.0032; Tukey’s post-hoc, JB-1 in sham versus Vx groups, *p*< 0.05). Similar effects were observed in the PAG (Fig. 4D; *F*_(2,18)_= 4.333, *p*= 0.0291; Tukey’s multiple comparison test, Vx JB-1 mice versus mice sham saline mice, *p*> 0.05) and the LC (Fig. 4E; *F*_(2,18)_= 19.37, *p*< 0.00001; Tukey’s post-hoc, JB-1 in sham versus Vx groups, *p*< 0.001).

**Figure 4.**
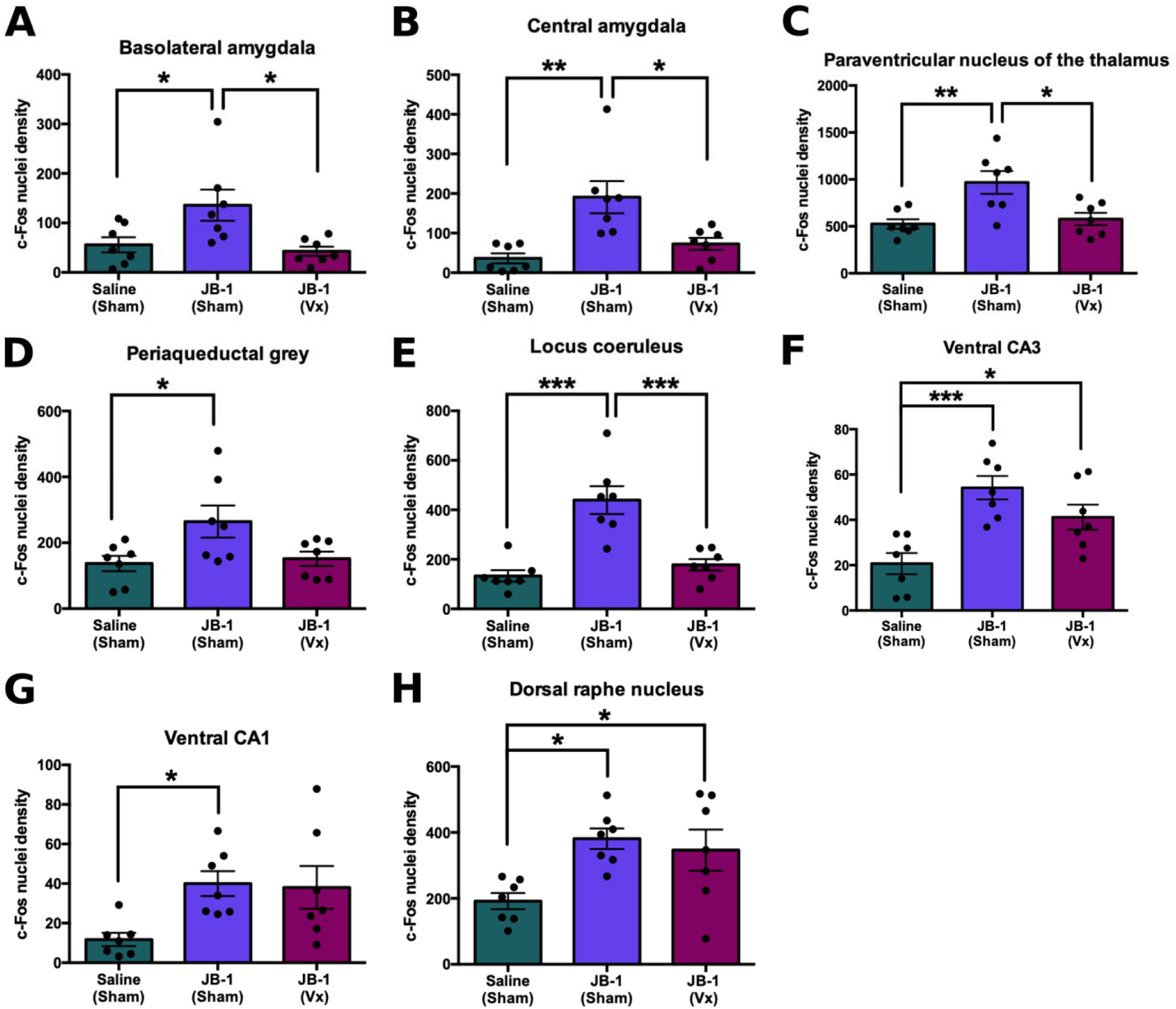
Effect of vagotomy on JB-1-induced c-Fos expression. Mean ± SEM c-Fos^+^ cell density (cells/mm^2^), quantified 165 minutes following administration of saline or live JB-1 to mice that underwent sham surgery or vagotomy (n=7 for all groups and regions). In all regions **(A-H)**, there was an overall main effect of treatment. Tukey post-hoc comparisons revealed that in most regions **(A-E)**, severing the vagus abolished JB-1-induced c-Fos expression. In certain regions however, severing the vagus did not affect c-Fos levels: in the vCA3 **(F)** and DRN **(H)**. In both regions, c-Fos levels in vagotomised animals administered the live bacteria were significantly greater than sham surgery mice administered saline, and no significantly different from sham surgery mice administered the live bacteria. * p<0.05, ** p<0.01, ***p<0.001.

In contrast, severing the vagus did not prevent an increase in c-Fos^+^ nuclei density in the vCA3 (Fig. 4F; *F*_(2,18)_= 10.80, *p*= 0.0008; Tukey’s post-hoc, JB-1 in sham versus Vx groups, *p*> 0.05; Vx and sham JB-1 mice versus sham saline mice, *p*< 0.05) and in the ventral CA1 (Fig. 4G; *F*_(2,18)_= 4.493, *p*= 0.0261), although post-hoc comparisons in the vCA1 failed to show a significant difference between Vx mice administered the live bacteria and sham surgery mice administered saline. In the DRN, both sham and Vx groups administered the live bacteria showed increased c-Fos immunoreactivity (Fig. 4H; *F*_(2,18)_= 5.604, *p*= 0.0128; Tukey’s multiple comparison test, JB-1 in sham versus Vx groups, *p*> 0.05, Vx and sham JB-1 mice versus sham saline mice, *p*< 0.05). These observations suggest that while signals to certain regions are vagal-dependent, other regions may also receive signals via vagal-independent pathways.

### 3.4. Chronic but not acute bacteria administration induces ΔFosB expression in distinct regions

ΔFosB is unique in its properties that allow it to accumulate in the brain in response to chronic stimulation, reflecting long-term neural adaptations (Nestler et al., 2001b; Mcclung et al., 2004). Since chronic JB-1 treatment engenders changes in time spent immobile (Fig. 5A; t_(20)_= 2.161, *p*= 0.0430) (Bravo et al., 2011; McVey Neufeld et al., 2018)—a marker for despair-like behaviour (Cryan et al., 2005), we investigated ΔFosB expression following acute versus chronic bacteria treatment.

**Figure 5.**
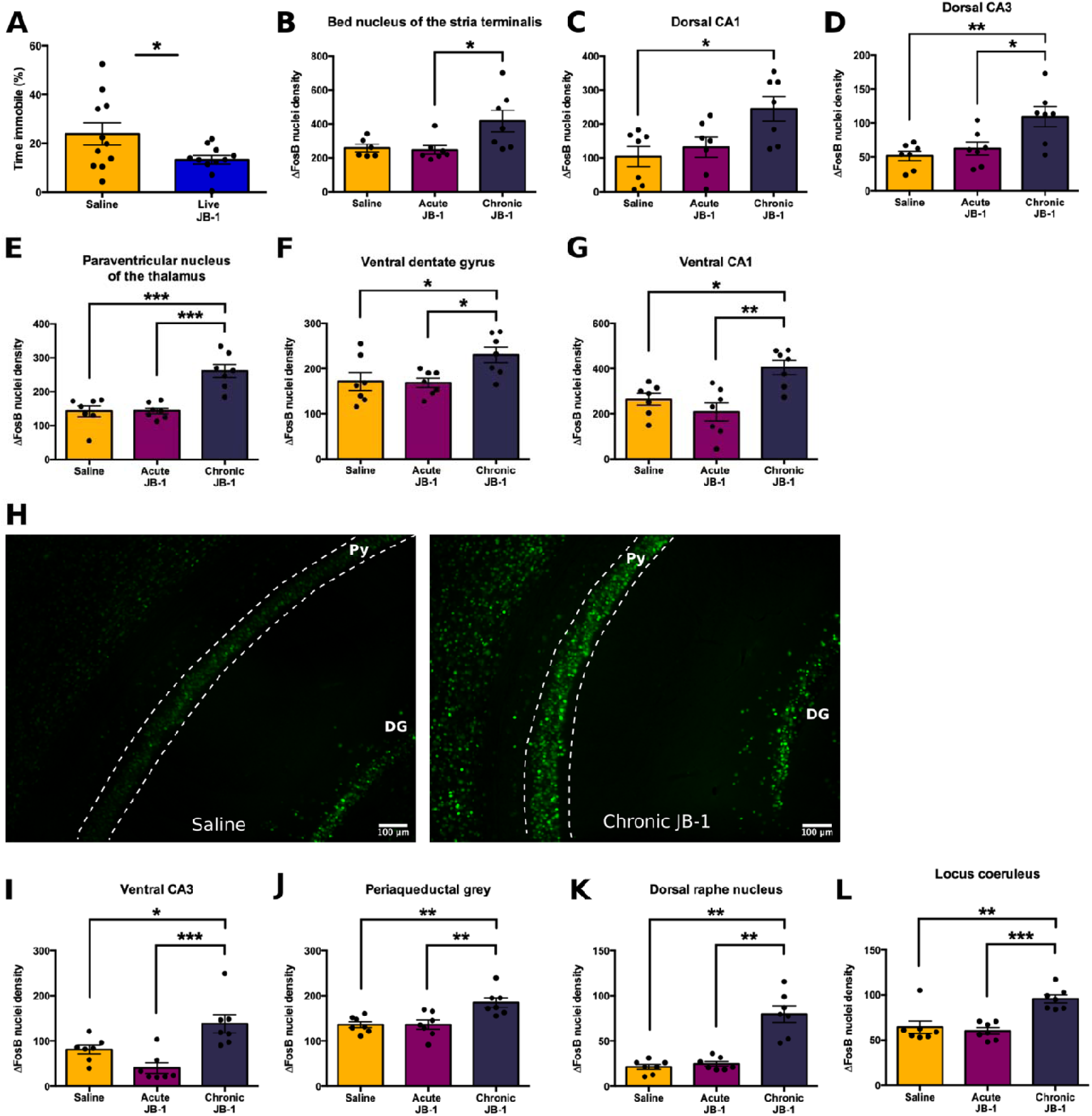
Chronic administration of JB-1 over two weeks induces ΔFosB expression in specific regions. **(A)** Percentage of time spent immobile during a six-minute tail suspension test, measured following chronic treatment with live bacteria. **(B-G, I-K)** ΔFosB^+^ cell density (cells/mm^2^), quantified 20-24h following two weeks of treatment. n=7 for all groups, with the exception of the BNST of the saline control group (n=6) **(B)**, due to damage incurred during dissection and tissue processing). **(H)** Representative images of ΔFosB staining in the vHpc * p<0.05, ** p<0.01, ***p<0.001. DG: dentate gyrus; Py: pyramidal cell layer, CA1.

Chronic administration induced ΔFosB expression in distinct brain regions, unlike acute JB-1 and saline treatments. In the BNST, there was a main effect of treatment (Fig. 5B; BNST, *H*= 7.311, *p*= 0.0259). Post-hoc comparisons revealed significantly greater ΔFosB^+^ nuclei density in chronic treated mice (Dunn’s multiple comparison test, *p*<0.05). Similar effects were observed in the dorsal CA1 and CA3 (Fig. 5C: dCA1, *F*_(2,18)_= 5.463, *p*= 0.0140; Fig. 5D: dCA3, *F*_(2,18)_= 7.786, *p*= 0.0037). Chronically treated mice exhibited greater ΔFosB levels relative to the saline treatment group in the dCA1 (*p*< 0.05), and versus acute (*p*< 0.05) and saline treatment groups (*p*< 0.01) in the dCA3. In the PVT, significantly greater ΔFosB levels were observed following chronic treatment (Fig. 5E: PVT, *F*_(2,18)_= 19.35, *p*< 0.0001; Tukey’s multiple comparison test, *p*< 0.001 vs acute and saline treatment groups). In the vHpc, increased ΔFosB expression was observed in the ventral DG (Fig. 5F: vDG, *F*_(2,18)_= 4.633, *p*= 0.0238; Tukey post-hoc, *p*< 0.05 vs acute and saline treatment), vCA1 (Fig. 5G & 5H: vCA1, *F*_(2,18)_= 9.452, *p*= 0.0016; Tukey post-hoc, *p*< 0.01 vs acute treatment and *p*< 0.05 vs saline treatment), and vCA3 (Fig. 5I: vCA3, *F*_(2,18)_= 11.21, *p*= 0.0007; Tukey post-hoc, *p*< 0.001 vs acute treatment and *p*< 0.05 vs saline treatment). Finally, similar effects of chronic treatment were observed on ΔFosB levels in the PAG (Fig. 5J: PAG, *F*_(2,18)_= 8.881, *p*= 0.0021; Tukey post-hoc, *p*< 0.01 vs acute and saline treatment), the DRN (Fig. 5K: DRN, *H*= 13.45, *p*= 0.0012; Dunn’s multiple comparison test, *p*< 0.01 vs acute and saline treatment), and the LC (Fig. 5L: LC, *F*_(2,18)_= 13.78, *p*= 0.0002; Tukey post-hoc, *p*< 0.001 vs acute treatment and *p*< 0.01 vs saline treatment).

In contrast, there was no effect of bacteria treatment on ΔFosB^+^ nuclei density in the NAc (Fig. 6A; *F*_(2,18)_= 2.811, *p*= 0.0866); PVH (Fig. 6B; *F*_(2,16)_= 2.537, *p*= 0.1104); dorsal DG (Fig. 6C; dDG, *F*_(2,18)_= 2.548, *p*= 0.1061); BLA (Fig. 6D; *F*_(2,17)_= 2.864, *p*= 0.0848); and CeA (Fig. 6E; *F*_(2,17)_= 2.945, *p*= 0.7487).

**Figure 6.**
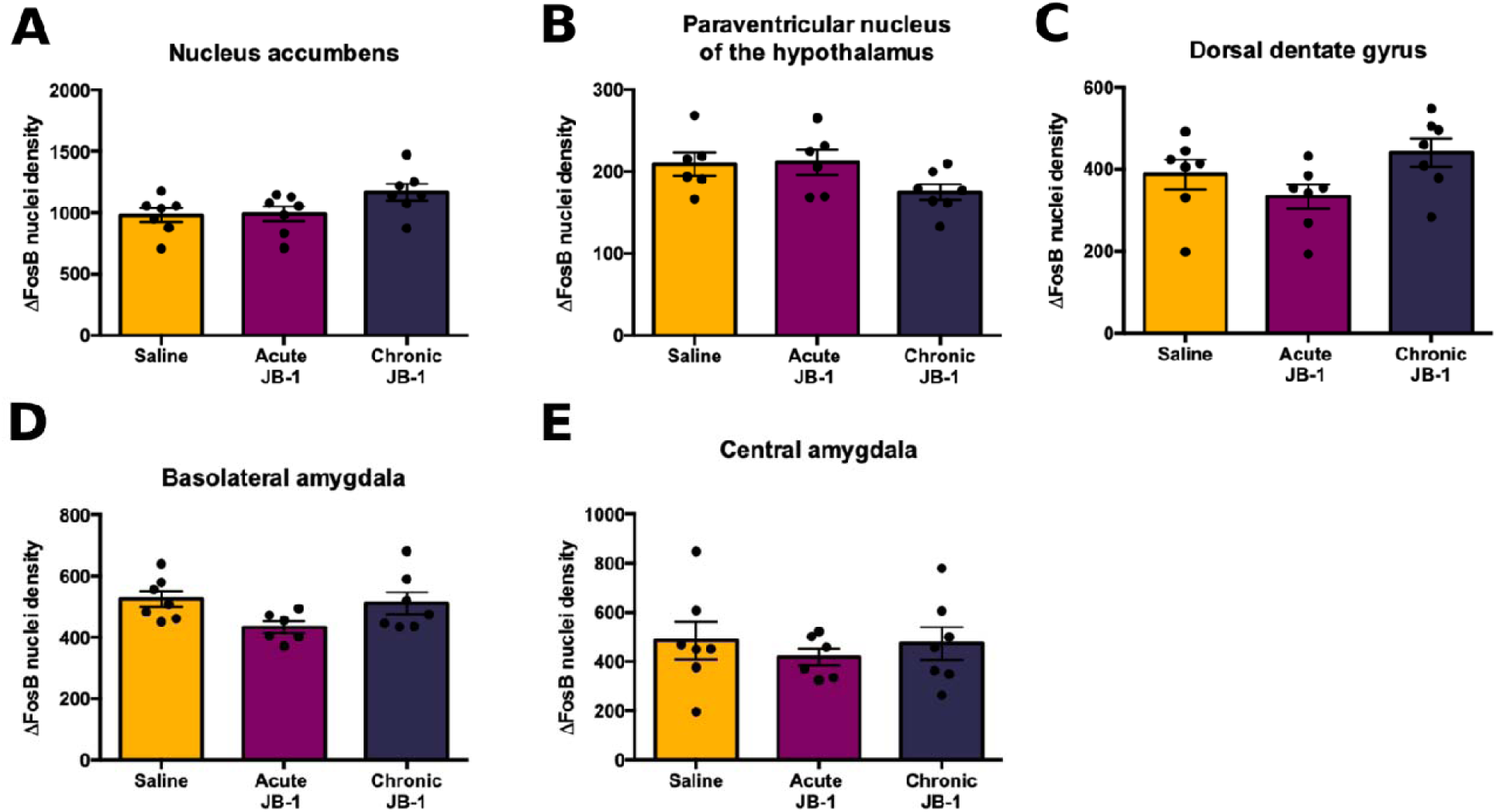
Chronic administration of JB-1 over two weeks does not affect ΔFosB expression in specific regions. **(A-E)** ΔFosB ^+^ cell density (cells/mm^2^), quantified 20-24h following two weeks of treatment. n=7 for all groups, with the exception of the PVH of the saline and acute treatment group **(B)**, and the BLA **(D)** and CeA **(E)** of the acute treatment group (n=6), due to damage incurred during dissection and tissue processing. There were no significant differences in ΔFosB levels between groups.

These data demonstrate that long-term oral treatment is necessary to induce ΔFosB expression in distinct regions throughout the brain.

## 4. Discussion

There exist multiple lines of evidence demonstrating the influence of intestinal bacteria via a proposed ‘gut-brain axis’ (Collins et al., 2012; Cryan and Dinan, 2012; Forsythe et al., 2016). However, little is known about the brain regions that are recruited by this form of signalling, the changes in the response pattern following acute versus chronic exposure to such signals, and the pathways that mediate these interactions. Here we show that oral administration of a specific bacterial strain induces expression of Fos gene products throughout the brain, within 165 minutes, and that the specific expression pattern differ between acute and chronic treatment. Although both live and heat-killed bacteria promote vagal firing, the effects on neuronal activity are mostly limited to the live bacteria. Additionally, severing the vagus nerve prevented c-Fos expression in many but not all brain regions, suggesting a critical role for the vagus but indicating the presence of additional signalling pathways.

Within 165 minutes, oral administration of the live bacteria induced c-Fos in the BLA, CeA, PVT, vCA1, vCA3, PAG, DRN, and LC (Fig. 1). Administration of heat-killed bacteria only increased c-Fos expression in the PVT. We chose the 165-minute time-point given that transit time to the mouse stomach and small intestine is ∼1 hour (Padmanabhan et al., 2013; Wehner et al., 2014), both of which are heavily innervated by the vagus nerve (Wang and Powley, 2000), and that c-Fos immunoreactivity in the brain peaks between 1-2 hours (Nestler et al., 2001b; Jung et al., 2014). While the data herein does not necessarily suggest that all of these distributed areas are integral to the effects of the bacteria, it aligns with previous works outlining regions that receive direct or indirect projections from the vagus nerve (Groves and Brown, 2005; Han et al., 2018), and those that are affected by vagal nerve stimulation (VNS)—an approved therapy in the treatment of refractory depression (Naritoku et al., 1995; Chae et al., 2003; Nemeroff et al., 2006; Cunningham et al., 2008; Furmaga et al., 2012; Aaronson et al., 2017). This is particularly relevant given the critical role of the vagus in mediating the behavioural effects of this particular bacterial strain (Bravo et al., 2011; Perez-Burgos et al., 2014). These observations are further underlined by studies showing that these regions also play a role in stress and mood-related processes, including the vHpc (Bagot et al., 2015; Padilla-coreano et al., 2016; Anacker et al., 2018; Jimenez et al., 2018), PVT (Beas et al., 2018), and LC (Mccall et al., 2017). Surprisingly, in the acute treatment experiments we did not observe increased c-Fos expression in the NTS, which is innervated by fibres from the nodose ganglia (Foley and DuBois, 1937; Nemeroff et al., 2006). This may have been a result of the chosen timepoint, or due to saline-induced gut distention, which also increases vagal firing (Perez-Burgos et al., 2013) and, consequently, c-Fos expression in the NTS (Schwarz et al., 2010; Jung et al., 2014), thus obscuring differences between treatment groups; or perhaps, cell groups in the NTS that respond to gut-derived vagal fibres are located more caudally in the brainstem (Schwarz et al., 2010; Han et al., 2018).

Severing vagal fibres below the diaphragm abolished JB-1-induced c-Fos expression in the BLA, CeA, PVT, PAG, and LC, while sparing the vHpc and DRN (Fig. 4). While we have shown that sub-diaphragmatic vagotomy abolishes the effects of JB-1 on anxiety-and depression-like behaviours (Bravo et al., 2011), it was unclear if this would also abolish the c-Fos response, or whether there exist additional pathways that transmit bacteria-derived signals. The data here demonstrate that vagotomy abolishes the c-Fos response in certain brain regions, while others continue to respond in the absence of an intact vagus nerve. c-Fos expression was no longer observed in regions that receive vagal afferents and are also activated by VNS: BLA, CeA, PVT, and LC (Naritoku et al., 1995; Chae et al., 2003; Cunningham et al., 2008; Han et al., 2018). In contrast, the DRN and vHpc exhibit elevated c-Fos levels following treatment in vagotomised mice. While both regions receive visceral afferents from the NTS, either through direct projections or via a multi-synaptic pathway (Peyron et al., 1996; Berthoud and Neuhuber, 2000; Castle et al., 2005), neither exhibits a c-Fos response to VNS as well (Naritoku et al., 1995; Cunningham et al., 2008). This further implicates a role for vagal-independent pathways in mediating JB-1-related signals to these regions, at least in acute settings. Putative pathways include spinal fibres, the activity of which can be modulated by different bacterial strains (Kamiya et al., 2006; Perez-Burgos et al., 2015). Additional pathways include soluble mediators such as cytokines, short-chain fatty acids, and bacteria- and host-derived metabolites (Forsythe et al., 2016). Bacteria also shed microvesicles, which contain a variety of molecules, including DNA and RNA (Dorward et al., 1989). In the case of JB-1, its microvesicles replicate the immune and enteric effects, resulting in increased levels of regulatory T cells and IL-10^+^ dendritic cells, as well as increased firing of intrinsic primary afferent neurons (Al-Nedawi et al., 2014). Future work should examine which of these pathways are readily recruited within the timeframe described in this study.

Chronic but not acute treatment increased ΔFosB expression in distinct regions throughout the brain (Fig. 5). Although previous studies have demonstrated induction of ΔFosB in response to acute stimulation (Nestler et al., 2001b; Mcclung et al., 2004; Perrotti et al., 2004), we did not observe a response to a single bacterial dose. Of the regions characterized by greater ΔFosB levels, some also exhibited robust c-Fos levels following acute treatment (Fig. 1), while others demonstrated no such response (Fig. 2: BNST, dorsal CA1 and CA3). Conversely, chronic treatment had no effect on ΔFosB levels in the BLA and CeA, which previously demonstrated treatment-induced c-Fos expression, suggesting that despite an acute response, not all regions undergo long-term adaptations to chronic treatment. The targets of FosB proteins have been studied extensively for their role in models of chronic drug use and mood disorders, including regulation of AMPA and NMDA glutamate receptor subunits (Mcclung et al., 2004; Vialou et al., 2010), NF-κB (Ang et al., 2001), dynorphin (Zachariou et al., 2006), and Ca^2+^/calmodulin-dependent protein kinase II (Robison et al., 2013). While all of the responsive regions may not be integral to the behavioural effects of the bacteria, the stability of ΔFosB uniquely position it as a potential mediator of neural changes, and its accumulating levels may thus reflect the presence of long-term adaptations (Mcclung et al., 2004; Nestler, 2015). Therefore, determining the distinct targets of ΔFosB in different brain regions may be critical to understanding its role in the behavioural changes associated with gut-brain signalling.

Within minutes of application to a jejunal segment, live and heat-killed bacteria increased the firing frequency of vagal fibres (Fig. 3). This change is driven by an increase in the firing of individual fibres rather than an increase in the number of active fibres (Mao et al., 2013; Perez-Burgos et al., 2013, 2014). Given that both live and heat-killed bacteria facilitate vagal firing but only the former induces widespread neuronal activity suggests intrinsic differences in signal encoding. This aligns with our data at 165 minutes following oral treatment, where we observed elevated firing frequency of vagal fibres only in mice treated with the live bacteria (Fig. 3D).It is possible these differences in signal encoding exist in the form and pattern of the elicited spike trains (Furness et al., 1999), or via additional signals provided by complementary pathways.

This study demonstrates that the CNS responds rapidly to a single oral administration of a specific bacteria. The widespread response is limited to the live bacteria, and there is a differential pattern of Fos expression following acute versus chronic administration. Severing the vagus prevents the acute response to bacterial treatment in certain regions that receive vagal projections. Given that vagus-mediated signalling is critical for the effects of JB-1 (Bravo et al., 2011) and other bacteria (Bercik et al., 2011) on brain chemistry and behaviour, some of the brain regions that are no longer responsive following vagotomy may play a critical role in mediating the effects of the bacteria. Based on these data, we also propose that signals from gut bacteria are relayed by additional, vagal-independent pathways along the gut-brain axis, thus enabling rich interactions between the periphery and the CNS. Future research will need to describe the precise bacteria-derived signals, as well as the gene targets of ΔFosB in order to understand the molecular underpinnings of gut-brain signalling.

## 5. Funding & Acknowledgments

The authors would like to thank Dr. Andrew Stanisz at the McMaster Brain-Body Institute for the preparation of bacterial treatments. This research was funded by a grant from the Office of Naval Research (N00014-14-1-0787), a CIHR CGS-D award to A.B. (GSD-148222), and an MD/PhD award to A.B. from the Research Institute of St. Joe’s Hamilton.

